# Chemogenetic enhancement of cAMP signaling renders hippocampal synaptic plasticity resilient to the impact of acute sleep deprivation

**DOI:** 10.1101/2022.03.11.483982

**Authors:** Emily Nicole Walsh, Mahesh Shivarama Shetty, Kamran Diba, Ted Abel

## Abstract

Sleep facilitates memory storage and even brief periods of sleep loss lead to impairments in memory, particularly memories that are hippocampus dependent. In previous studies, we have shown that the deficit in memory seen after sleep loss is accompanied by deficits in synaptic plasticity. Our previous work has also found that sleep deprivation is associated with reduced levels of cyclic adenosine monophosphate (cAMP) in the hippocampus, and that the reduction of cAMP mediates the diminished memory performance. Based on these findings, we hypothesized that cAMP acts as a mediator for not only the cognitive deficits caused by sleep deprivation, but also the observed deficits in synaptic plasticity. In this study, we expressed the heterologous *Drosophila melanogaster* Gαs-protein coupled octopamine receptor (DmOctβ1R) in mouse hippocampal neurons. This receptor is selectively activated by the systemically injected ligand (octopamine), thus allowing us to increase cAMP levels in hippocampal neurons during a five-hour sleep deprivation period. Our results show that chemogenetic enhancement of cAMP during the period of sleep deprivation prevents deficits in a persistent form of long-term potentiation (LTP) that is induced at the Schaffer collateral synapses in the hippocampal CA1 region. We also found that elevating cAMP levels only in the early or later half of sleep deprivation successfully prevented LTP deficits. These findings reveal that cAMP-dependent signaling pathways are key mediators of sleep deprivation at the synaptic level. Targeting these pathways could be useful in designing strategies to prevent the impact of sleep loss.

**Significance statement:** Insufficient sleep is an issue with significant health and socioeconomic implications. This includes a negative impact on memory consolidation. Previous studies in mice found that acute sleep deprivation leads to deficits in hippocampal synaptic plasticity and memory, which are associated with reduced levels of the signaling molecule cAMP. In this study, we used a chemogenetic strategy to enhance cAMP levels in specific hippocampal neurons during sleep deprivation. We found that this made synaptic plasticity resilient to the negative effects of sleep deprivation. These findings reveal that cAMP-dependent signaling pathways are key mediators of sleep deprivation and that targeting these pathways could be useful in designing strategies to prevent the impact of sleep loss.

## Introduction

It has been known for many years that sleep is an important part of long-term memory storage. Even short periods of sleep can improve declarative memory (Tucker et al., 2006; Van Der Helm et al., 2011), and loss of sleep leads to impairments in memory in both mice and humans (Graves et al., 2003; Palchykova et al., 2006; Rasch & Born, 2013). One way that sleep is believed to carry out this function is by altering the synaptic plasticity that underlies memory consolidation (Yang & Gan, 2012; Yang et al., 2014; Li et al., 2017). This is particularly true in the hippocampus, an important brain region for the consolidation of episodic memories, which undergoes changes in molecular and cellular signaling, and demonstrates impaired synaptic plasticity in response to sleep deprivation (SD). A single, brief period of SD by gentle handling impairs hippocampal protein synthesis (Tudor et al., 2016; Lyons et al., 2020), and hippocampal cyclic adenosine monophosphate (cAMP) signaling— the latter of which leads to reduced activation of downstream effectors such as cAMP response element binding protein (CREB) (Vecsey et al. 2009), LIM kinase (LIMK) and cofilin (Havekes et al., 2016b). These effectors have important roles in synaptic plasticity. The alterations in molecular signaling caused by SD lead to selective deficits in persistent forms of long-term potentiation (LTP) that have been detected at the Schaffer collateral synapses in the stratum radiatum of hippocampal area CA1 (Vecsey et al., 2009; Vecsey et al., 2018; Wong et al., 2019). Multiple studies have shown that persistent forms of LTP, which last through the late phase of LTP (L-LTP), are dependent on cAMP signaling, transcription, and translation of new proteins (Frey et al., 1988; Frey et al., 1993; Nguyen et al., 1994; Lynch, 2004).

The reduction in cAMP levels in SD has been attributed, at least in part, to an increase in the protein levels of cAMP-specific phosphodiesterase PDE4A5 (Vecsey et al., 2009). Previous work has also shown that the SD-induced deficits in persistent forms of LTP could be rescued by treating hippocampal slices with a PDE4 inhibitor to prevent cAMP degradation (Vecsey et al., 2009). A similar strategy of inhibiting PDEs to prevent cAMP degradation also prevents the hippocampal memory deficits that are associated with acute SD (Vecsey et al. 2009; Heckman et al., 2020). While these observations collectively point to cAMP signaling as a key pathway in mediating the effects of sleep on synaptic plasticity, previous studies were unable to modify cAMP levels in a spatiotemporally precise manner. These limitations can be addressed by chemogenetic approaches which involve expression of a heterologous receptor in a specific brain region or cell-type and its selective activation by controlled administration of its ligand. One such approach utilizes the expression of the heterologous *Drosophila melanogaster* Gαs-protein coupled octopamine receptor (DmOctβ1R), which when activated by the delivery of its ligand octopamine, leads to enhancements in cAMP levels (Balfanz et al., 2005; Isiegas et al., 2008). We previously utilized this approach to virally express the DmOctβ1R in hippocampal CaMKIIα neurons, and selectively activated the receptors by systemically delivering octopamine at specific time points during the acute SD period. The resultant elevation of cAMP levels in hippocampal neurons during acute SD prevented the associated deficit in hippocampus-dependent memory (Havekes et al., 2014). In the current study, we use the same approch of virus-mediated expression of the DmOctβ1R in hippocampal neurons and its activation by systemic delivery of octopamine during SD to investigate whether elevating hippocampal cAMP levels during SD would also provide resilience to hippocampal synaptic plasticity. Our results show that increasing cAMP levels in hippocampal neurons during SD prevents the deficit in a form of long-lasting LTP in the CA1 region. Together, our findings demonstrate the role of cAMP in the deterioration of hippocampal functions following acute SD and indicate that cAMP signaling is a key mediator of the impact of SD.

## Materials and Methods

### Subjects

Male C57BL/6J mice (Jackson Laboratory #000664), 3-4 months of age, were used for all the experiments. Prior to the start of the experiment, mice were group housed (up to 5 per cage) in soft bedding cages. Food (NIH-31 irradiated modified mouse diet #7913) and water were provided *ad libitum*. These mice were on a 12:12 light schedule, lights-on at 8 AM. The start of the lights-on period marks zeitgeber time zero (ZT0). For mice that underwent surgery, the surgery was performed between 10-12 weeks of age. Mice were maintained in group housing during the recovery from surgery and during the weeks preceding the start of the experiment. Experiments were conducted according to National Institutes of Health guidelines for animal care and use and were approved by the Institutional Animal Care and Use Committee (IACUC) at the University of Iowa.

### Viral vectors and surgeries

To manipulate cAMP levels in hippocampal neurons we used an AAV construct (serotype 9; AAV9-CaMKIIα0.4-DmOctβ1R -HA (titer 1.24E+14 genome copies (GC)/mL)) containing the *Drosophila melanogaster* octopamine receptor type 1β (DmOctβ1R) (Balfanz, Strünker, Frings, & Baumann, 2005), which when bound by the octopamine ligand, activates adenylyl cyclase and stimulates cAMP production. The plasmid was generated using Geneart and packaged by the University of Pennsylvania Viral Vector Core. The construct is expressed under the CaMKIIα promoter (0.4Kb fragment) and also contains an HA tag to facilitate visualizing the expression of the receptor. The stock virus was diluted to a lower titer (1.24E+13 GC/mL) in saline solution (0.9% sodium chloride, Hospira Inc.) prior to infusion into the hippocampus. To assess the effects of octopamine in the absence of the receptor, an enhanced green fluorescent protein (eGFP) under the CaMKIIα promoter was used (AAV9-CaMKIIα-eGFP-WPRE; Addgene #50469, 2.4E+13 GC/mL). We infused the AAV in the dorsal hippocampus through the following coordinates relative to bregma: anteroposterior (AP) −1.9 mm, mediolateral (ML) ±1.5 mm, dorsoventral (DV) −1.5 mm. Mice were induced and maintained anaesthetized with isoflurane for the surgery. The AAV suspension was infused bilaterally (1000 nl per hemisphere at a rate of 200 nl/min) using a NanoFil syringe (World Precision Instruments, NanoFil 10 ul) through a 33G beveled needle (World Precision Instruments, # NF33BV-2), controlled by a micro-syringe pump (World Precision Instruments, Microinjection Syringe Pump, # UMP3T-2). We allowed 4 weeks to pass between the initial virus delivery and the day of the experiment to allow for the virus to express.

### Drug preparation

1 mg of (±)-Octopamine hydrochloride (Millipore Sigma # O0250-5G) was dissolved in 2 ml of 0.9% saline solution, to obtain 0.5 mg/ml concentration of octopamine solution. This solution was prepared fresh on the day of the experiment. The volume was administered based on the weight of the mouse at a dose of 1 mg/kg. For vehicle controls, the appropriate volume of 0.9% saline was used for injections. Octopamine solution or saline was administered to mice by intraperitoneal (i.p.) injection.

## Experimental design

### Sleep deprivation

All mice were singly housed seven days prior to the sleep deprivation (SD) or non-sleep deprivation (NSD) day. Each cage had corncob bedding (Envigo, Teklad ¼” corncob, #7907), and a small amount of soft bedding for mice to make an adequate nest. These cages were equipped with water bottles and wire hoppers to hold food, and mice had *ad libitum* access to food and water at all times, including during SD. Mice were handled for 5 days prior to the experiment by the same researcher conducting the experiments. This allowed the mice to habituate to the experimenter, room, and the tapping stimulation that was used on the cage to keep mice in the SD group awake. For handling, the mice were taken to the SD room, and each mouse was held in the experimenter‘s palm for 2 minutes. They were then placed back in their cages, and cages were tapped for 2 minutes to habituate them to the stimuli that would be used in the gentle handling method of sleep deprivation. For mice in an experiment requiring injection, injection habituation started 2-3 days prior to the day of SD (0.9% saline vehicle, 0.1ml injection i.p.). The injection was administered after handling, and prior to mice being returned to their cage for 2 minutes of tapping. SD began at ZT0 and continued for 5 hours using the gentle handling method (Hagewoud et al., 2010; Prince et al., 2014; Vecsey et al., 2009) in which taps to the cage were administered as needed to keep mice awake. When taps were no longer sufficient the mice received a “cage shake” which was a motion of the cage to offset the balance of the mouse and rouse them. Mice in the NSD group were housed and handled identically, but on the day of the experiment were instead kept in their behavioral colony housing room throughout the 5-hour period.

### Slice electrophysiology

Immediately after the SD or NSD period (at the end of ZT5), mice were cervically dislocated and the hippocampi were rapidly dissected in artificial cerebrospinal fluid (aCSF; NaCl 124 mM, KCl 4.4 mM, MgSO_4_.7H_2_O 1.3 mM, CaCl_2_.2H_2_O 2.5 mM, NaH_2_PO_4_.H_2_O 1 mM, NaHCO_3_ 26.2 mM, D-Glucose 10 mM, PH ∼7.4, Osmolarity ∼300 mOsm) with continuous flow of carbogen (95% oxygen, 5% carbon dioxide). 400 µm-thick transverse hippocampal slices were prepared from the dorsal 2/3 portion of both the hippocampi by a manual McIlwain slicer (Stoelting), as previously described (Shetty et al. 2015). The slices were placed in a netted interface chamber (Fine Science Tools, Foster City, CA) and incubated at 28°C for at least 2 hours in oxygenated aCSF (perfused at 1 mL/min) before starting electrophysiological recordings. For all recordings, a monopolar, lacquer coated stainless-steel electrode (A-M Systems #571000) was positioned in the CA1 stratum radiatum to stimulate Schaffer collaterals, and an aCSF-filled glass electrode (2–5 MΩ resistance) was also placed in the CA1 stratum radiatum to record field excitatory postsynaptic potentials (fEPSPs). For test stimulation, a biphasic, constant current pulse (100 μs duration per phase) was delivered using an isolated pulse stimulator (Model 2100, A-M Systems, Carlsborg, WA) and recorded using IE250 Intracellular Electrometer (Warner Instruments). Data were low-pass filtered at 2 kHz (LPF100B, Warner Instruments) and acquired at 20 kHz using pClamp 10 software and Axon Digidata 1440/1550 digitizers (Molecular Devices, Union City, CA). For every slice, an input-output curve (stimulation intensity vs fEPSP amplitude) was generated, and the baseline stimulation intensity was set to elicit ∼40% of the maximal fEPSP amplitude. In all the experiments, test stimulation was performed once every minute, including for 20 minutes to establish a stable baseline prior to long-term potentiation (LTP) induction. LTP was induced with a spaced 4-train stimulation paradigm (four 100 Hz, 1 s trains delivered 5 minutes apart, at the baseline intensity) and recordings were continued for 160 minutes. The data were analyzed using Clampfit 10 analysis software (Axon Molecular Devices, Union City, CA). In every experiment, the fEPSP initial slopes were normalized to the 20-minute baseline average and expressed as percentages. Input–output characteristics were assessed by quantifying fEPSP and presynaptic fiber volley (PFV) amplitudes in response to increasing stimulus intensity (0 to 70 µA, at 5 µA increments). Paired-pulse facilitation (PPF) was assessed by delivering two pulses at baseline intensity at different inter-pulse intervals (300, 200, 100, 50 and 25 ms). Facilitation was quantified by the ratio of the second fEPSP amplitude to the first. In all the electrophysiology experiments, reported “n” values refer to the number of mice, and data from replicate slices from the same mouse are averaged. Mean and standard error of the mean (SEM) are reported in figure legends and results.

### Immunohistochemistry (IHC)

To confirm the viral expression of DmOctβ1R-HA in hippocampal tissue, we used an anti-HA IHC protocol with chromogenic 3,3′-Diaminobenzidine (DAB) staining. Mice were perfused with 1X PBS (∼10 mL), followed by 4% PFA (∼10 mL). Brains were extracted into 4% PFA solution and left overnight (4°C) before being transferred to 30% sucrose solution (4°C). Once equilibrated, the brains were sliced into 30 µm thick sections on a cryostat (Leica 3050S). All steps for DAB staining were done with gentle rotation and at room temperature unless otherwise stated. First, slices were washed in 1X PBS three times for 5 minutes each, then incubated for 25 minutes in H_2_O_2_ (0.3% H_2_O_2_ in 1X PBS). Next, the slices were washed with 1X PBS for 30 minutes before being preincubated with 5% normal goat serum and 0.1% Triton-X in 1X PBS. Following preincubation, slices were incubated overnight with the primary antibody (HA-Tag (C29F4), Cell Signaling #3724S) 1:100 in 1X PBS plus 0.1% Triton-X and 1% normal goat serum. The next day, slices were washed for 3×10 minutes with 1X PBS before being incubated for 5 hrs with the secondary antibody (Biotinylated, Vector lab, cat no. BA-1000) 1:500 in 1X PBS and 1% normal goat serum. Afterwards, they were washed 3×10 min with 1X PBS, then incubated for 2hr with ABC kit (VECTASTAIN® Elite® ABC HRP Kit (Peroxidase, Standard), cat no. K-6100; 1:100 of both components in 1X PBS). The slices were then washed for 4hr in 1X PBS (one 15 min wash in 1XPBS, then again with fresh 1XPBS x3 every 1-1.5hr), before being moved into the DAB (DAB-HCL, Electron Microscopy Sciences, Fisher Scientific catalog #50-980-352) solution (0.15 mg DAB/mL of 1X PBS, add 100 μL 0.1% H_2_O_2_ to every 5mL of the solution right before starting the stain). Slices are incubated in the DAB solution for 8 minutes. After the DAB step, the slices were washed 3×10 min in 1X PBS to arrest further reaction. Slices were moved out of 1X PBS onto Superfrost Plus (Fisherbrand) slides, allowed to dry overnight, and then coverslipped with Permount (Fisher Chemical™ Permount™ Mounting Medium, Fisher Scientific # SP15-100).

## Statistical analysis

Power analyses were performed for all experiments at an alpha of 0.05 and a desired power level of 0.80 to estimate the number of mice needed. The estimated effect sizes were based on previous publications using similar methods. All the surgeries and sleep deprivation were performed by one experimenter and the electrophysiology experiments were performed by another experimenter blind to the identity of the mice or condition. Experiments from respective control and experimental mice were performed side-by-side on any given day. In the LTP experiments, the average fEPSP slopes over the course of the recording are expressed as percentages of the respective baseline average in each group. Electrophysiology data were extracted using Clampfit 10 (Axon Molecular Devices) and Statistical analyses were performed using GraphPad Prism 9. Data were tested for normality and the maintenance of LTP was assessed by comparing the average fEPSP slopes from the last 20 min of the recording using two-tailed unpaired t-tests. Input-output and PPF data were compared using two-way repeated measures ANOVA. For all analyses the statistical significance was set at *p* < 0.05. In figures, * refers to a *p* value < 0.05, ** refers to *p* < 0.01.

## Results

We have previously shown that 5 hours of SD by the gentle handling method leads to deficits in long-lasting LTP induced by spaced tetanic-train stimulation (Vecsey et al., 2009; Wong et al., 2019). Given the differences in animal housing and minor changes in the electrophysiological rig compared with the earlier study, in the first series of experiments we confirmed the impact of acute SD on spaced 4-train LTP. Mice were either sleep deprived using gentle handling from ZT0-5 or allowed to sleep (non-sleep deprived; NSD) for the same duration. At ZT5, hippocampal slices were prepared for spaced 4-train LTP recordings at the Schaffer collateral synapses in the CA1 stratum radiatum (**Fig. 1A**). The data showed clear deficits in the persistence of this long-lasting form of LTP in the SD group, where the potentiation decayed to baseline levels within 2 hours, compared to stable long-lasting LTP in the NSD group (**Fig. 1B**). The maintenance of LTP was assessed by comparing the mean potentiation over the last 20 minutes of the recording between the two conditions. The mean potentiation of the NSD group (229 ± 29.71%) and the SD group (86.29 ± 7.72%) revealed a significant deficit in the SD group (**Fig. 1C**; two-tailed unpaired t-test, t(9) = 4.250, *p* = 0.002, η^2^= 0.664). These data confirm previous observations (Vecsey et al., 2009) that a brief period of SD leads to impairments in long-lasting LTP induced by spaced 4-train stimulation.

**Figure 1.**
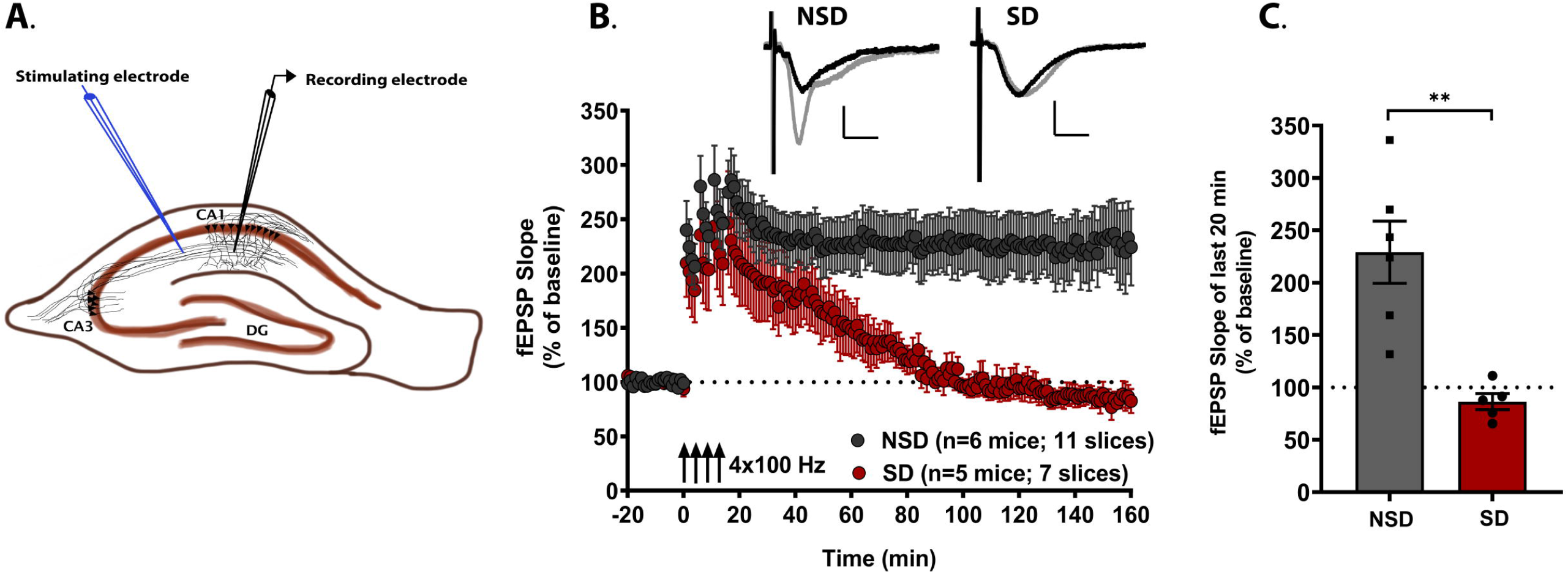
Brief sleep deprivation impairs the persistence of cAMP-PKA signaling-dependent LTP in the hippocampal CA1 region. **(A)** Schematic representation of a transverse hippocampal slice showing the positioning of stimulating and recording electrodes in the CA1 stratum radiatum. **(B)** Long-lasting LTP induced by spaced-4-train stimulation protocol (four 100 Hz, 1 s stimulation trains, spaced at 5 minutes) is impaired in slices from mice subjected to 5 h of sleep deprivation (SD) compared to non-sleep deprived (NSD) mice. The representative fEPSP traces shown for each group are sampled at baseline (black trace) and at the end of the recording (grey trace). Scale bars for traces: 2 mV vertical, 5 ms horizontal. **(C)** Persistence of LTP, assessed by comparing the mean potentiation over the last 20 minutes of the recording between the NSD group (229 ± 29.71%) and SD group (86.29 ± 7.72%), shows a significant deficit in the SD group (two-tailed t-test, t(9) = 4.250, p = 0.002, η2 = 0.6640).

Next, we investigated the effect of chemogenetically enhancing cAMP levels in hippocampal neurons during the SD period. We virally expressed (**Fig. 2A**) the Gαs-coupled DmOctβ1R or eGFP control in the hippocampal neurons of adult mice under the CaMKIIα promoter. We confirmed the expression of DmOctβ1R by DAB staining using an antibody against the HA tag on the receptor, which showed clear hippocampus-restricted expression of the receptor (**Fig. 2B**). Mice expressing either DmOctβ1R or eGFP virus were subjected to five hours of SD from ZT0-ZT5. During SD, mice received two intraperitoneal (i.p.) injections of either octopamine (1 mg/kg) or saline vehicle. These injections were administered at the start (ZT0) and halfway through (ZT2.5) the SD period (**Fig. 2C**). This timing and dosage were based on our previous study, which showed effective elevation of hippocampal cAMP levels using these conditions (Havekes et al., 2014). At the end of the SD period, hippocampal slices were prepared and LTP was induced by the spaced 4-train protocol. We then compared the time-course of LTP between the octopamine and saline groups within each viral condition (**Fig. 2D, 2F**).

**Figure 2.**
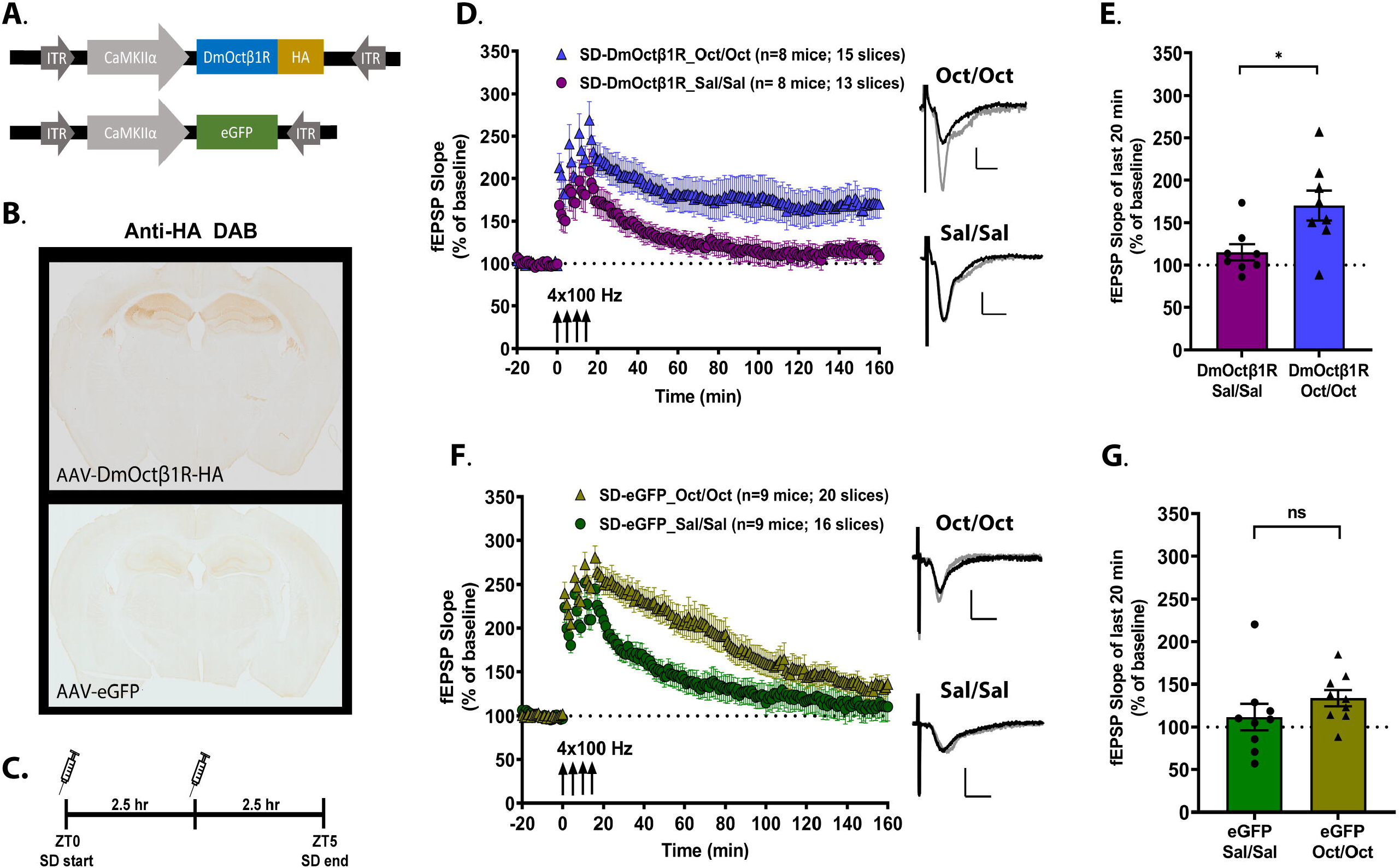
Chemogenetic enhancement of cAMP signaling during acute sleep deprivation confers resilience to the associated deficits in spaced 4-train LTP. **(A)** Schematic representation of the AAV constructs used to drive the expression of *Drosophila melanogaster* octopamine receptor (DmOctβ1R) or eGFP in hippocampal CaMKIIα expressing neurons. **(B)** Representative images of coronal brain sections from mice expressing DmOctβ1R-HA, or control eGFP, which were probed by chromogenic DAB staining against the HA tag. The upper panel shows the restricted expression of DmOctβ1R in the hippocampal subregions, and the lower panel shows the lack of signal from the eGFP expressing slice. **(C)** A schematic representation of the timeline of injections during the course of 5 h sleep deprivation (SD), starting at ZT0. Mice, expressing either DmOctβ1R or eGFP in the hippocampus, receive two injections of either the ligand octopamine or saline at ZT0 and at ZT2.5. At the end of the 5 h SD, hippocampi are sliced and LTP induced by spaced 4-train LTP is investigated. **(D)** Following 5 h SD, mice expressing DmOctβ1R and receiving two injections of octopamine (Oct/Oct) show no deficits in the long-lasting spaced-4-train LTP whereas those receiving two injections of saline (Sal/Sal) show impairments. **(E)** Persistence of LTP in the DmOctβ1R-expressing mice, assessed by comparing the mean potentiation over the last 20 min of recordings between the saline vehicle group (115 ± 9.64%) and octopamine group (170.5 ± 17.74%), shows significantly enhanced potentiation in the octopamine group (two-tailed unpaired t-test, t(14) = 2.749, *p =* 0.016, η^2^ = 0.351). **(F)** In mice expressing eGFP, spaced 4-train LTP persistence is impaired in both the octopamine (Oct/Oct) and saline (Sal/Sal) injected conditions. **(G)** The mean potentiation over the last 20 min of recordings between the saline group (111.7 ± 15.61%) and octopamine group (134 ± 9.53%) showed no significant difference between the two groups (two-tailed unpaired t-test, t(16) = 1.219, *p =* 0.24, η^2^ = 0.085). The representative fEPSP trace for each group shown in **D** and **F** was sampled at the respective baseline (black trace) and at the end of the recording (grey trace). Scale bars for traces: 2 mV vertical, 5 ms horizontal.

In SD mice expressing DmOctβ1R and receiving two injections of saline vehicle, LTP decayed to baseline levels within two hours (**Fig. 2D**), similar to the non-injected wildtype SD mice in Fig.1B. In contrast, in SD mice expressing the DmOctβ1R that received two octopamine injections, LTP was persistent for the duration of the recording (**Fig. 2D**). The persistence of LTP was evaluated by comparing the mean potentiation over the last 20 min of recordings between the DmOctβ1R saline group (115 ± 9.64%) and the DmOctβ1R octopamine group (170.5 ± 17.74%), and revealed significantly enhanced potentiation in the octopamine group (**Fig. 2E**; two-tailed unpaired t-test, t(14) = 2.746, *p* = 0.016, η^2^ = 0.351). We also assessed basal synaptic transmission by comparing the input-output responses (stimulation intensity versus fEPSP or PFV amplitude) and paired-pulse facilitation (PPF), a very short-term form of plasticity, and found no significant differences between the two groups in any of these measures (**Supplemental Fig. S1A, S1B, S1C**). These results show that enhancing cAMP signaling in hippocampal neurons during SD confers resilience against the negative impact of brief SD on persistent synaptic plasticity.

To confirm that the resilience of LTP observed in the DmOctβ1R-expressing mice injected with octopamine was due to the activation of the heterologous receptor, we investigated the effect of two injections of octopamine or saline during SD (at ZT0 and ZT2.5) in mice with virally expressed eGFP in hippocampal neurons. When we compared basal synaptic transmission measures, we observed some differences between the saline and octopamine SD eGFP groups in the fEPSP amplitudes (**Supplemental Fig. 2A**) and PFV amplitudes (**Supplemental Fig. 2B**), but no significant difference in PPF (**Supplemental Fig. 2C**). eGFP-expressing SD mice showed deficits in the maintenance of LTP following spaced 4-train LTP induction, regardless of whether they received octopamine or saline injections (**Fig. 2F**). The mean potentiation over the last 20 min of the recordings in the SD eGFP saline group (111.7 ± 15.61%) and SD eGFP octopamine group (134 ± 9.53%) showed no significant difference (**Fig. 2G**; two-tailed unpaired t-test, t(16) = 1.219, *p* = 0.240, η^2^ = 0.085). Although we observed some differences in the time-course of LTP decay between these groups, the persistence of LTP, which is the hallmark of long-lasting synaptic plasticity (Frey et al., 1993), was impaired in both conditions. Overall, these findings demonstrate that the persistent LTP in slices from DmOctβ1R-expressing mice that received octopamine injections was due to effects on the heterologous Gαs-receptor that enhance intracellular cAMP levels.

Our findings (**Fig. 2**) demonstrate that chemogenetically enhancing cAMP levels at multiple time points during SD can render hippocampal LTP resilient to the impact of 5 hours of acute SD. Other studies have shown that shorter windows of sleep deprivation can also produce alterations in gene expression in the hippocampus (Delorme et al., 2021) and the cortex (Cirelli & Tononi, 2000) and impair memory and synaptic plasticity (Prince et al., 2014). These findings raise a question about whether the enhancement of cAMP levels in the early or late windows of SD have consistent or different effects on the maintenance of LTP. To examine the possibility that a single injection of octopamine during either the early or later half of the 5h SD period could sufficiently prevent deficits in LTP, we performed another set of experiments using mice virally expressing DmOctβ1R in hippocampal neurons. The design was similar to the octopamine experiments above, except mice in one group now received the injection of octopamine only at ZT0 and saline at ZT2.5 (Oct/Sal), while mice in the other group received saline at ZT0 and octopamine at ZT2.5 (Sal/Oct) (**Fig.3A**). At the end of the five-hour SD, hippocampal slices were prepared for spaced 4-train LTP recordings in the CA1 stratum radiatum. Interestingly, persistent LTP was observed for both of these conditions (**Fig.3B**). Comparing the mean potentiation over the last 20 min of recordings, we found no significant difference between the Oct/Sal group (230.6 ± 16.87%) and the Sal/Oct group (182.8 ± 19.27%), although the octopamine injection in the early window (Oct/Sal) appeared to trend towards being more effective (**Fig.3C**; two-tailed unpaired t-test, t(15) = 1.844, *p* = 0.085, η^2^ = 0.185). We also assessed the basal synaptic transmission and PPF and found no significant differences between the two groups in any of these measures (**Supplemental Fig. 3A-C**). These results suggest that activating cAMP signaling either in the first or second half of the SD period can prevent the decay of LTP induced by sleep loss, supporting the notion that the impact of SD on synaptic plasticity builds over time.

**Figure 3.**
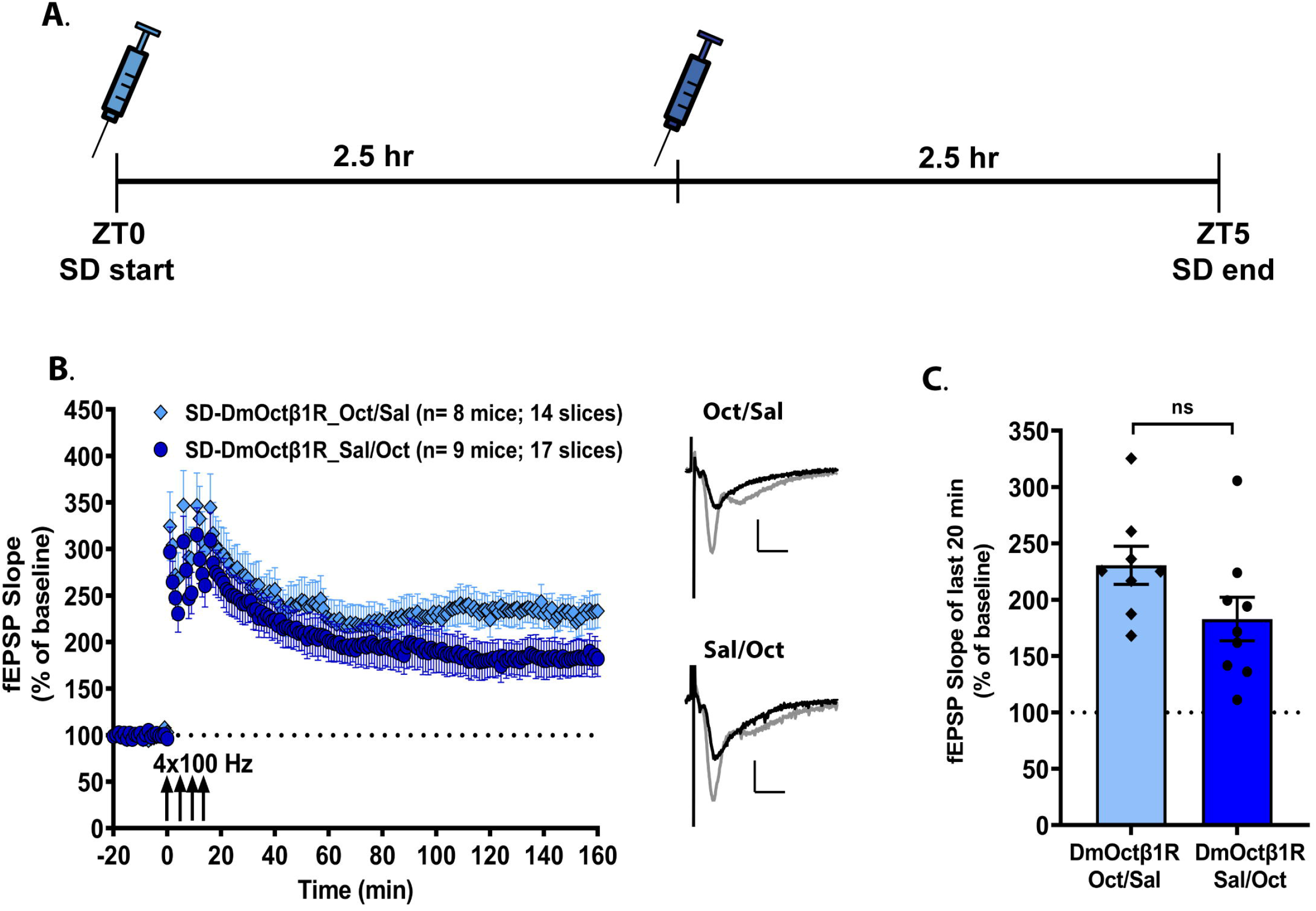
Enhancement of hippocampal cAMP signaling, either early or late during brief sleep deprivation, effectively prevents the associated deficits in long-lasting LTP. **(A)** A schematic representation of the timeline of injections during the course of 5 h sleep deprivation. Mice expressing virally DmOctβ1R in hippocampal neurons receive an injection of octopamine at either ZT0 or at ZT2.5. Mice that receive octopamine injection at ZT0 get a saline injection at ZT2.5 (Oct/Sal), and mice that get octopamine injection at ZT2.5 receive a saline injection at ZT0 (Sal/Oct). **(B)** A single injection of octopamine to chemogenetically enhance cAMP signaling either during the early or later periods of 5 h SD is effective in preventing the impact of SD on spaced 4-train LTP. The representative fEPSP traces shown for each group are sampled at baseline (black trace) and at the end of the recording (grey trace). Scale bars for traces: 2 mV vertical, 5 ms horizontal. **(C)** The mean potentiation over the last 20 min of recordings between the Oct/Sal group (230.6 ± 16.87%) and the Sal/Oct group (182.8 ± 19.27%) shows no significant difference between the conditions, although the injection in the early window (Oct/Sal) trends towards being more effective (two-tailed unpaired t-test, t(15) = 1.844, *p =* 0.085, η^2^ = 0.185).

## Discussion

There is growing evidence that the disruption of cAMP signaling is responsible for impairments in hippocampus-dependent processes following acute SD (Vecsey et al., 2009; Havekes et al., 2014; Wong et al., 2019). Sleep deprivation causes a decrease in cAMP levels in the hippocampus (Vecsey et al., 2009), which may be driven by increased levels and activity of the phosphodiesterase PDE4A5 (Vecsey et al., 2009; Wong et al., 2019). PDE4A5 overexpression in hippocampal neurons mimics the memory and plasticity phenotype of SD (Havekes, et al., 2016a), and blockade of PDE4A5 activity by overexpression of a catalytically inactive form of PDE4A5 prevents the memory deficits that follow SD (Havekes et al., 2016b). There is also a reduction in the overall activity of protein kinase A (PKA) in SD, and thus a decrease in the phosphorylation of important downstream effectors (Wong et al., 2019). The present study was designed to investigate if increasing cAMP levels in hippocampal neurons during SD is enough to protect hippocampal plasticity from alterations in molecular signaling caused by SD.

Our data demonstrate that increasing cAMP levels during SD has a protective effect for synaptic plasticity. We found that activating the heterologous Gαs-coupled DmOctβ1 receptor with the ligand octopamine prevented the decay of spaced 4-train LTP, which is a form of long-lasting LTP known to be dependent on cAMP-PKA signaling, transcription, and translation (Frey et al., 1988; Frey et al., 1993; Nguyen et al., 1994; Huang & Kandel, 1994; Abel et al., 1997; Malleret et al., 2001). Spaced 4-train LTP is vulnerable to acute SD, as was reported previously (Vecsey et al., 2009), and confirmed in our experiments. Mice without chemogenetic elevation of hippocampal cAMP (either eGFP expressing conditions, or DmOctβ1R with saline) show vulnerability in this form of LTP following SD, suggesting that the resilience we observed was mediated by the activation of the heterologous receptor and not due to an off-target effect of octopamine. Although the eGFP-expressing mice that received two octopamine injections showed deficits in the persistence of LTP following SD, the rate of LTP decay showed some differences compared to eGFP-expressing mice receiving saline. This intriguing difference was accompanied by an altered basal synaptic transmission, and together, these observations hint at some possible effects of octopamine that are not mediated by the heterologous receptor. A potential explanation for these effects could be the action of octopamine on trace amine-associated receptors (TAARs) in the hippocampus, which may then modulate neurotransmitter signaling (Berry, 2004; Wolinski et al., 2007; Zucchi et al., 2009). The relatively low expression of TAARs in the hippocampus (Borowsky et al., 2001; Lindemann et al., 2008) and the requirement of high micromolar concentrations of octopamine to effect other neurotransmitter signaling (Berry, 2004), suggest however that this is unlikely to be the case. With the decay of the eGFP-saline LTP back to baseline it demonstrates that if octopamine is having an effect on TAARs, that effect is not substantial enough to confer resilience against the impact of SD on LTP. Nevertheless, our results act as a guidepost for future studies using similar approaches to consider such possibilities.

Our results with the single injection of octopamine either in earlier or later half of SD period show that LTP is made resilient by elevated cAMP levels regardless of the timepoint within the course of SD that it occurred. In our experiments, we administered octopamine at the beginning of SD or in the middle of the 5-hour deprivation period and found that either injection prevented the deficit in plasticity. Furthermore, there was no significant difference in the effectiveness of the early or late injection. This is interesting, because evidence suggests that the effects of acute SD build over time (Marks and Wayner, 2005). Three hours of SD cause mild impairments in LTP in the dentate gyrus of the hippocampus, but effects are more severe with six-hour SD (Marks and Wayner, 2005). Our present and previous work attributes the SD-related dysfunction in the hippocampus to changes in cAMP levels. It is possible that the disruption of cAMP-dependent molecular signaling pathways develops progressively over the SD period and that preventing the disruption anywhere along the course prevents it from falling below a threshold to impact LTP persistence. From this perspective, injection of octopamine either early or later during SD leads to lesser disruption in cAMP signaling than would normally occur in a full 5 hour span of SD. The similar effectiveness of either an early or late injection of octopamine suggests that enhancing intracellular cAMP at either time point prevented the disruption in cAMP associated with SD from reaching the critical threshold needed to cause a deficit in LTP. It is important to note that the full time-course of cAMP degradation during SD remains unknown. That we did not see a difference between the early or late injection of octopamine raises new questions about the hierarchy and dynamics of disruption of molecular signaling events during sleep loss, which will be the goal of future experiments.

Future studies may focus on the different molecular cascades induced during the period of SD, as well as on the time course of cAMP degradation and PDE upregulation. It would be important to map the hierarchy or sequence of signaling events underlying the impact of SD. This would help identify key causal events that lead to the multitude of effects of SD on neuronal structure and function. It would also be interesting to investigate whether the chemogenetic strategy used in this study is also effective in conferring resilience to other forms of synaptic plasticity, such as theta-burst stimulation LTP, potentiation induced by forskolin, and synaptic tagging, which are all impacted by brief SD (Vecsey et al. 2009; Vecsey et al. 2018). Additionally, investigating pre-or postsynaptic compartment-specific mechanisms impacted by SD would be exciting and can be facilitated by manipulations restricted to specific hippocampal subregions. Finally, given that there are changes structural plasticity with SD, such as changes in dendritic spine density in specific hippocampal regions (Raven et al., 2018; Bolsius et al., 2021), future studies could also address whether elevation in hippocampal cAMP levels also confers resilience against these changes. These studies would provide more information about how the deficits of SD emerge and how they can be prevented.

This work provides important insights on the hippocampal mechanisms that can be altered to provide resilience to the impact of acute sleep loss. Greater understanding of this central mechanism can also point to possible strategies for intervention in diseases with sleep loss, by detailing the mechanisms necessary to prevent the decline in cognitive processing that accompany it.

## Acknowledgements

**(Including funding sources)** The study was supported by the National Institutes of Health R01 Grant MH117964 to T.A and K.D, the National Institutes of Health R21 Grant MH117788 to K.D, and the University of Iowa Hawkeye Intellectual and Developmental Disability Research Center (P50 HD 103556; L. Strathearn and TA, multi PIs). TA is the Roy J. Carver Chair of Neuroscience.

## Supplemental Figures

**Supplemental Figure 1.**
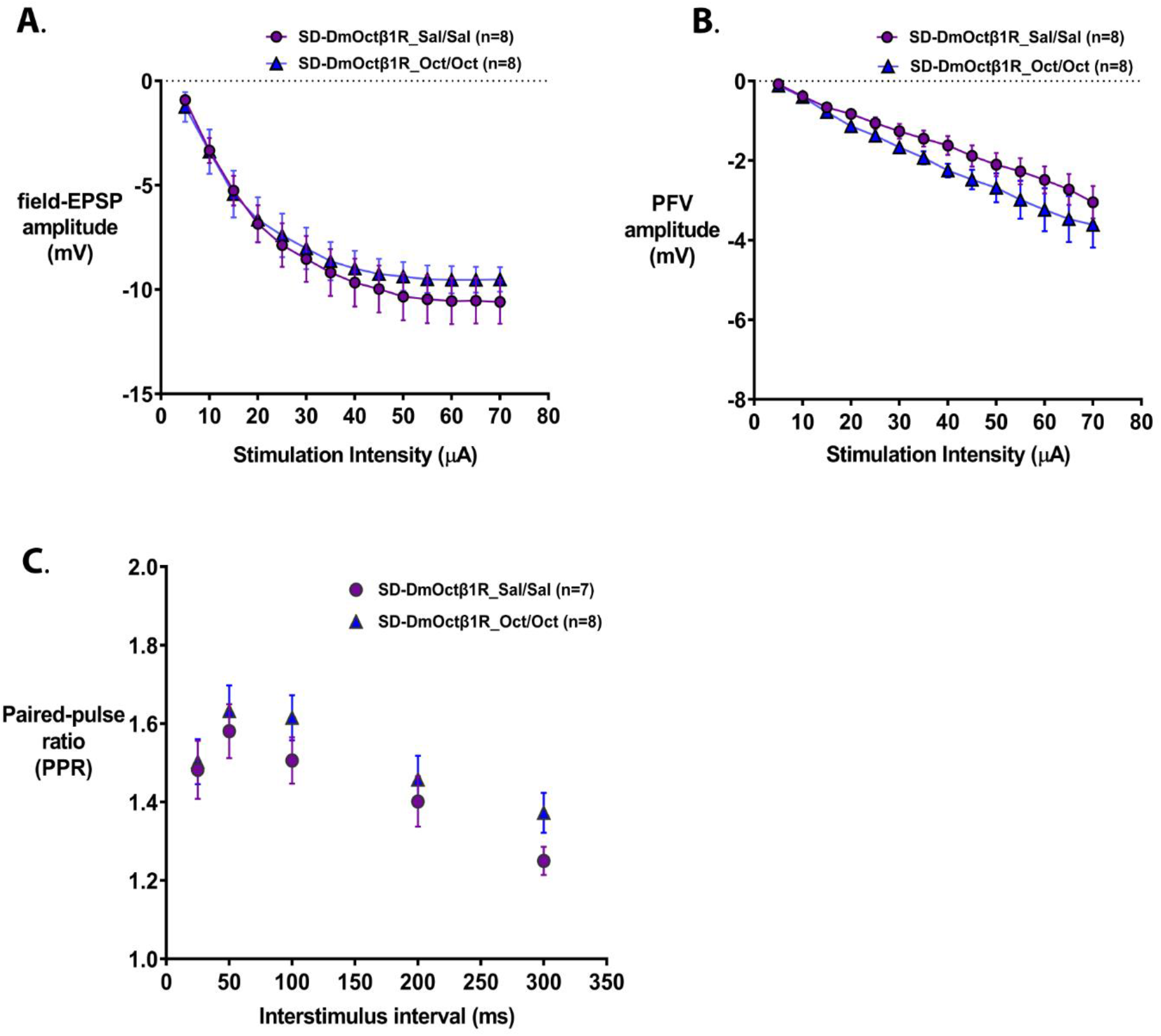
Basal synaptic transmission and paired-pulse facilitation in mice virally expressing DmOctβ1R and receiving two injections of octopamine or saline during sleep deprivation. **(A)** Basal field-EPSP amplitudes are not significantly different between the saline group and the octopamine group (Two-way repeated measures ANOVA; F (1,14) = 0.202; *p* = 0.660). **(B)** Presynaptic fiber volley (PFV) amplitudes are not significantly different between the saline group and the octopamine group (Two-way repeated measures ANOVA; F (1,14) = 2.113; *p* = 0.168). **(C)** Paired-pulse facilitation over a range of interstimulus intervals is not significantly different between the saline group and the octopamine group (Two-way repeated measures ANOVA; F (1,13) = 0.825; *p* = 0.380).

**Supplemental Figure 2.**
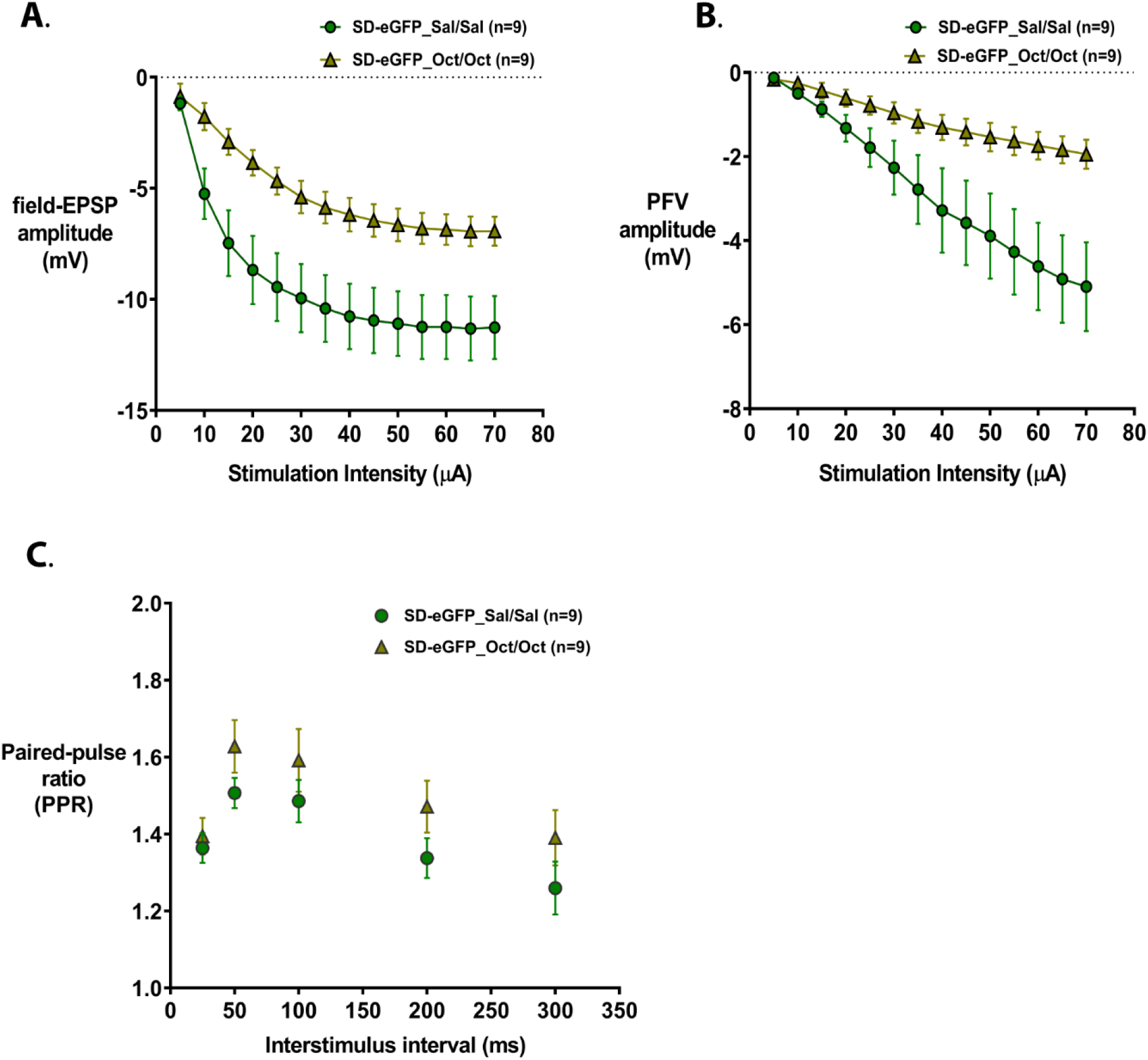
Basal synaptic transmission and paired-pulse facilitation in mice virally expressing eGFP and receiving octopamine or saline injections during sleep deprivation. **(A)** Basal field-EPSP amplitudes showed significant difference between the saline group and the octopamine group (Two-way repeated measures ANOVA; F (1,16) = 8.224; *p* = 0.011). **(B)** Presynaptic fiber volley (PFV) amplitudes showed significant difference between the saline group and the octopamine group (Two-way repeated measures ANOVA; F (1,16) = 5.371; *p* = 0.034). **(C)** Paired-pulse facilitation over a range of interstimulus intervals is not significantly different between the saline group and the octopamine group (Two-way repeated measures ANOVA; F (1,16) = 2.079; *p* = 0.169).

**Supplemental Figure 3.**
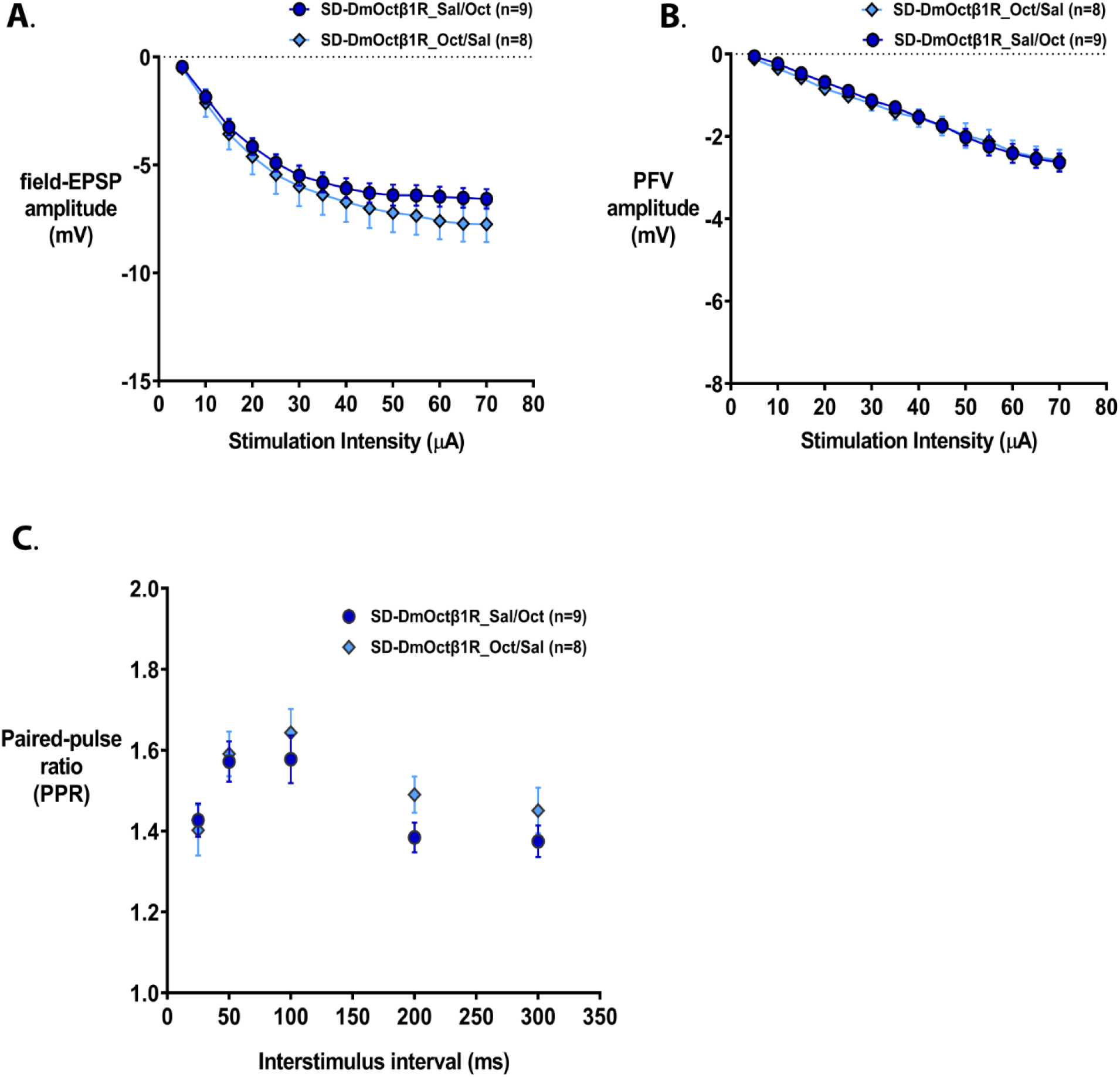
Basal synaptic transmission and paired-pulse facilitation in mice virally expressing DmOctβ1R and receiving a single octopamine injection (either at ZT0 or at ZT2.5) during sleep deprivation. **(A)** Basal field-EPSP amplitudes are not significantly different between the saline group and the octopamine group (Two-way repeated measures ANOVA; F (1,15) = 0.628; *p* = 0.440). **(B)** Presynaptic fiber volley (PFV) amplitudes are not significantly different between the saline group and the octopamine group (Two-way repeated measures ANOVA; F (1,15) = 0.026; *p* = 0.874). **(C)** Paired-pulse facilitation over a range of interstimulus intervals is not significantly different between the saline group and the octopamine group (Two-way repeated measures ANOVA; F (1,15) = 0.715; *p* = 0.411).

## Notes

**Conflict of interest statement:** All the authors declare that there are no conflicts of interest.

### Competing Interest Statement

The authors have declared no competing interest.

## References

Abel T, Nguyen PV, Barad M, Deuel TA, Kandel ER, Bourtchouladze R (1997) Genetic demonstration of a role for PKA in the late phase of LTP and in hippocampus-based long-term memory. Cell 88(5):615–26.

Balfanz S, Strünker T, Frings S, Baumann A (2005) A family of octopamine [corrected] receptors that specifically induce cyclic AMP production or Ca2+ release in Drosophila melanogaster. J Neurochem 93(2):440–51. Erratum in: J Neurochem 2005.

Berry MD (2004) Mammalian central nervous system trace amines. Pharmacologic amphetamines, physiologic neuromodulators. J Neurochem 90(2):257–71.

Bolsius YG, Meerlo P, Kas MJ, Abel T, Havekes R (2022) Sleep deprivation reduces the density of individual spine subtypes in a branch-specific fashion in CA1 neurons. J Sleep Res 31(1):e13438.

Borowsky B, Adham N, Jones KA, Raddatz R, Artymyshyn R, Ogozalek KL, Durkin MM, Lakhlani PP, Bonini JA, Pathirana S, Boyle N, Pu X, Kouranova E, Lichtblau H, Ochoa FY, Branchek TA, Gerald C (2001) Trace amines: identification of a family of mammalian G protein-coupled receptors. Proc Natl Acad Sci U S A 98(16):8966–71.

Cirelli C, Tononi G (2000) Differential expression of plasticity-related genes in waking and sleep and their regulation by the noradrenergic system. J Neurosci 20(24):9187–94.

Delorme J, Wang L, Kodoth V, Wang Y, Ma J, Jiang S, Aton SJ (2021) Hippocampal neurons’ cytosolic and membrane-bound ribosomal transcript profiles are differentially regulated by learning and subsequent sleep. Proc Natl Acad Sci U S A 118(48):e2108534118.

Frey U, Huang YY, Kandel ER (1993) Effects of cAMP simulate a late stage of LTP in hippocampal CA1 neurons. Science 260(5114):1661–4.

Frey U, Krug M, Reymann KG, Matthies H (1988) Anisomycin, an inhibitor of protein synthesis, blocks late phases of LTP phenomena in the hippocampal CA1 region in vitro. Brain Res 452(1-2):57–65.

Graves LA, Heller EA, Pack AI, Abel T (2003) Sleep deprivation selectively impairs memory consolidation for contextual fear conditioning. Learn Mem 10(3):168–76.

Havekes R, Bruinenberg VM, Tudor JC, Ferri SL, Baumann A, Meerlo P, Abel T (2014) Transiently increasing cAMP levels selectively in hippocampal excitatory neurons during sleep deprivation prevents memory deficits caused by sleep loss. J Neurosci 34(47):15715–21.

Havekes R, Park AJ, Tolentino RE, Bruinenberg VM, Tudor JC, Lee Y, Hansen RT, Guercio LA, Linton E, Neves-Zaph SR, Meerlo P, Baillie GS, Houslay MD, Abel T (2016a) Compartmentalized PDE4A5 Signaling Impairs Hippocampal Synaptic Plasticity and Long-Term Memory. J Neurosci 36(34):8936–46.

Havekes R, Park AJ, Tudor JC, Luczak VG, Hansen RT, Ferri SL, Bruinenberg VM, Poplawski SG, Day JP, Aton SJ, Radwańska K, Meerlo P, Houslay MD, Baillie GS, Abel T (2016b) Sleep deprivation causes memory deficits by negatively impacting neuronal connectivity in hippocampal area CA1. Elife 5:e13424.

Heckman PRA, Roig Kuhn F, Meerlo P, Havekes R (2020) A brief period of sleep deprivation negatively impacts the acquisition, consolidation, and retrieval of object-location memories. Neurobiol Learn Mem 175:107326.

Huang YY, Kandel ER (1994) Recruitment of long-lasting and protein kinase A-dependent long-term potentiation in the CA1 region of hippocampus requires repeated tetanization. Learn Mem 1(1):74–82.

Isiegas C, McDonough C, Huang T, Havekes R, Fabian S, Wu LJ, Xu H, Zhao MG, Kim JI, Lee YS, Lee HR, Ko HG, Lee N, Choi SL, Lee JS, Son H, Zhuo M, Kaang BK, Abel T (2008) A novel conditional genetic system reveals that increasing neuronal cAMP enhances memory and retrieval. J Neurosci 28(24):6220–30.

Li W, Ma L, Yang G, Gan WB. REM sleep selectively prunes and maintains new synapses in development and learning. Nat Neurosci. 2017 Mar;20(3):427–437. doi: 10.1038/nn.4479. Epub 2017 Jan 16. PMID: 28092659; PMCID: PMC5535798.

Lindemann L, Meyer CA, Jeanneau K, Bradaia A, Ozmen L, Bluethmann H, Bettler B, Wettstein JG, Borroni E, Moreau JL, Hoener MC (2008) Trace amine-associated receptor 1 modulates dopaminergic activity. J Pharmacol Exp Ther 324(3):948–56.

Lyons LC, Chatterjee S, Vanrobaeys Y, Gaine ME, Abel T (2020) Translational changes induced by acute sleep deprivation uncovered by TRAP-Seq. Mol Brain 13(1):165.

Malleret G, Haditsch U, Genoux D, Jones MW, Bliss TV, Vanhoose AM, Weitlauf C, Kandel ER, Winder DG, Mansuy IM (2001) Inducible and reversible enhancement of learning, memory, and long-term potentiation by genetic inhibition of calcineurin. Cell 104(5):675–86.

Marks CA, Wayner MJ (2005) Effects of sleep disruption on rat dentate granule cell LTP in vivo. Brain Res Bull 66(2):114–9.

Nguyen PV, Abel T, Kandel ER (1994) Requirement of a critical period of transcription for induction of a late phase of LTP. Science 265(5175):1104–7.

Palchykova S, Winsky-Sommerer R, Meerlo P, Dürr R, Tobler I (2006) Sleep deprivation impairs object recognition in mice. Neurobiol Learn Mem 85(3):263–71.

Prince TM, Wimmer M, Choi J, Havekes R, Aton S, Abel T (2014) Sleep deprivation during a specific 3-hour time window post-training impairs hippocampal synaptic plasticity and memory. Neurobiol Learn Mem 109:122–30.

Rasch B, Born J (2013) About sleep’s role in memory. Physiol Rev 93(2):681–766.

Raven F, Meerlo P, Van der Zee EA, Abel T, Havekes R (2019) A brief period of sleep deprivation causes spine loss in the dentate gyrus of mice. Neurobiol Learn Mem 160:83–90.

Tudor JC, Davis EJ, Peixoto L, Wimmer ME, van Tilborg E, Park AJ, Poplawski SG, Chung CW, Havekes R, Huang J, Gatti E, Pierre P, Abel T (2016) Sleep deprivation impairs memory by attenuating mTORC1-dependent protein synthesis. Sci Signal 9(425):ra41.

Tucker MA, Hirota Y, Wamsley EJ, Lau H, Chaklader A, Fishbein W (2006) A daytime nap containing solely non-REM sleep enhances declarative but not procedural memory. Neurobiol Learn Mem 86(2):241–7.

van der Helm E, Gujar N, Nishida M, Walker MP (2011) Sleep-dependent facilitation of episodic memory details. PLoS One 6(11):e27421.

